# Reovirus core proteins λ1 and σ2 promote stability of disassembly intermediates and influence early replication events

**DOI:** 10.1101/2020.03.18.997874

**Authors:** Stephanie Gummersheimer, Pranav Danthi

**Affiliations:** Department of Biology, Indiana University, Bloomington, Indiana, USA

## Abstract

The capsids of mammalian reovirus contain two concentric protein shells, the core and the outer capsid. The outer capsid is comprised of µ1-σ3 heterohexamers which surround the core. The core is comprised of λ1 decamers held in place by σ2. After entry into the endosome, σ3 is proteolytically degraded and µ1 is cleaved and exposed to form ISVPs. ISVPs undergo further conformational changes to form ISVP*s, resulting in the release of µ1 peptides which facilitate the penetration of the endosomal membrane to release transcriptionally active core particles into the cytoplasm. Previous work has identified regions or specific residues within reovirus outer capsid that impact the efficiency of cell entry. We examined the functions of the core proteins λ1 and σ2. We generated a reovirus T3D reassortant that carries strain T1L derived σ2 and λ1 proteins (T3D/T1L L3S2). This virus displays a lower ISVP stability and therefore converts to ISVP*s more readily. To identify the basis for lability of T3D/T1L L3S2, we screened for hyper-stable mutants of T3D/T1L L3S2 and identified three point mutations in µ1 that stabilize ISVPs. Two of these mutations are located in the C-terminal ϕ region of µ1, which has not previously been implicated in controlling ISVP stability. Independent from compromised ISVP stability, we also found that T3D/T1L L3S2 launches replication more efficiently and produces higher yields in infected cells. In addition to identifying a new role for the core proteins in disassembly events, these data highlight that core proteins may influence multiple stages of infection.

**IMPORTANCE:** Protein shells of viruses (capsids) have evolved to undergo specific changes to ensure the timely delivery of genetic material to host cells. The 2-layer capsid of reovirus provides a model system to study the interactions between capsid proteins and the changes they undergo during entry. We tested a virus in which the core proteins were derived from a different strain than the outer capsid. We found that this mismatched virus was less stable and completed conformational changes required for entry prematurely. Capsid stability was restored by introduction of specific changes to the outer capsid, indicating that an optimal fit between inner and outer shells maintains capsid function. Separate from this property, mismatch between these protein layers also impacted the capacity of virus to initiate infection and produce progeny. This study reveals new insights into the roles of capsid proteins and their multiple functions during viral replication.

## INTRODUCTION

In order to successfully launch replication, a virus must protect, transport and deliver its genome into the host cell. The viral capsid is a complex mechanical container with the primary function of assembling around the viral genome in one host cell, exiting that host cell and releasing the genome in another host cell. Therefore, the capsid has two demands that appear to be in direct conflict with one another. It must be stable enough to protect and transport the viral genome, but it also must be dynamic, poised to react to the right environment at the right time to perform functions during entry and to allow for replication of the genome. Viruses have evolved capsid proteins that are capable of these dynamic mechanical interactions in a number of diverse and fascinating ways. While some capsids are made up of only a single type of capsid protein, others have more complex combinations of proteins. One family of viruses, *Reoviridae*, have double or even triple layered capsids.

The *Reoviridae* family of viruses is made up of non-enveloped virions with capsids that are composed of 1 to 3 concentric protein shells that surround 9 to 12 ds RNA genome segments (1–3). While the outer layers of the multilayered capsids are proteolytically processed and undergo conformational changes during entry, the innermost capsid or core which contains the genome, remains intact throughout the remainder of replication (4). The core is a complex molecular machine that is capable of producing fully capped and functional RNA transcripts for translation by the host machinery (5, 6). The capsid proteins must, therefore, be capable of multiple functions in addition to their protective and structural roles. The outer capsid proteins must be poised to undergo the conformational changes required to release the cores and allow them to be transcriptionally active. While studies of mammalian reovirus have provided many insights into how outer capsid proteins regulate and mediate entry events that lead to these conformational changes, the role of the core proteins in cell entry events remains unclear.

The capsids of mammalian reovirus are made up of two concentric protein shells, the core and the outer capsid (4). The outer capsid is primarily made up of µ1-σ3 heterohexamers (7) (Fig 1). These surround the core which is comprised of λ1 decamers held in place by σ2 (8). Turrets, made up of λ2 pentamers, protrude from the core at the five-fold axis of symmetry. The σ1 attachment protein forms trimers that are anchored in the λ2 turret (8, 9). After entry, σ3 is proteolytically degraded within the endosome and µ1 is cleaved into µ1δ and µ1ϕ fragments (10, 11). These µ1 fragments remain particle associated at this stage and the particle is referred to as an infectious subvirion particle or ISVP (12–14). Further conformational changes within the endosome result in cleavage of µ1δ to form µ1N and δ (15, 16). These particles are no longer infectious and are referred to as ISVP*. Release of µ1 peptides (µ1N and ϕ) results in penetration of the endosomal membrane and the deposition of the transcriptionally active cores into the cytoplasm (13, 17, 18).

**Figure 1.**
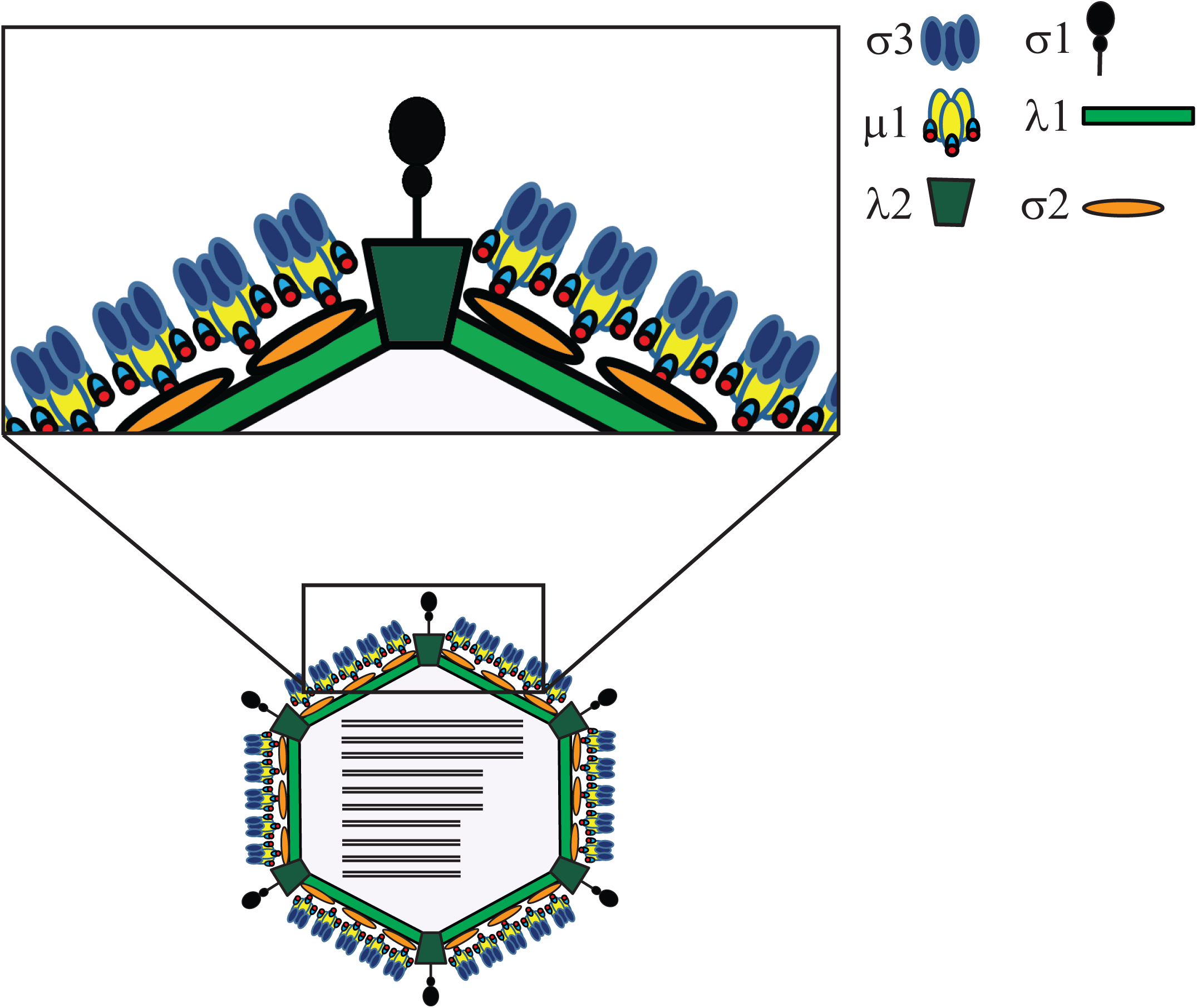
Schematic representation of reovirus capsid proteins.

Previous work in identifying interactions and proteins involved in reovirus disassembly and cell entry have focused on the outer capsid proteins. Here, we focused on the structural functions of the core proteins λ1 and σ2. We characterized a reassortant virus containing the λ1 encoding gene segment L3 and the σ2 encoding gene segment S2 (T3D/T1L L3S2). We found that this new virus displays a higher ISVP-to-ISVP* conversion efficiency or lower ISVP stability. We identified three point mutations in µ1 that increase the stability of T3D/T1L L3S2. Two of these mutations are located in the ϕ region of µ1, which has not previously been implicated in maintaining ISVP stability or controlling ISVP-to-ISVP* conversion. Additionally, this reassortant virus has increased growth resulting from higher transcription levels and, subsequently, higher protein production. This role does not appear to be related to the enhanced ISVP-to-ISVP* conversion of this virus. These results provide insights into understanding the structural functions of the core proteins and how their interactions may influence disassembly and early replication steps during infection.

## RESULTS

### ISVP to ISVP* conversion efficiency of T3D is altered by the T1L core proteins λ1 and σ2

Single gene reassortants between prototype reovirus strains T1L and T3D, which contain mismatches between outer capsid proteins, display altered capsid stability and efficiency of disassembly intermediate formation (19–21). To determine whether the core proteins λ1 and σ2 play a role in ISVP-to-ISVP* conversion efficiency, we characterized the properties of T3D/T1L L3S2. This virus contains the λ1-encoding gene segment L3 and the σ2-encoding gene segment S2 from T1L in an otherwise T3D genetic background. The resulting virus therefore contains major core proteins that do not match the outer capsid. The λ1 protein from T3D and T1L are 99.3% identical (9 amino acid differences out of 1,275) and the σ2 proteins are 98.8% identical (5 amino acid differences out of 418) (22–24). While σ2 interacts with the outer capsid µ1 trimer, λ1 is not known to interact with any outer capsid proteins (25).

To test overall virion stability, we incubated T3D and T3D/T1L L3S2 over a series of elevated temperatures and tested the protease stability of the major viral capsid proteins. Such an approach has been used previously to test stability of virions and viral entry intermediates (19). Based on the similarity in their protease sensitivity profiles, we think that virions of T3D/T1L L3S2 do not display significant changes in stability in comparison to the parent strain, T3D (Fig 2A). To determine if the mismatches in this reassortant virus alter stability of ISVPs or efficiency of ISVP-to-ISVP* conversion, we generated ISVPs of T3D and T3D/T1L L3S2 and incubated them over a gradient of increasing temperatures. ISVP* conversion was determined as a measure of trypsin sensitivity of the µ1 δ fragment (26). In comparison to the parent strain T3D, T3D/T1L L3 ISVPs have significantly reduced stability or enhanced ISVP-to-ISVP* conversion *in vitro* (Fig 2B).

**Figure 2.**
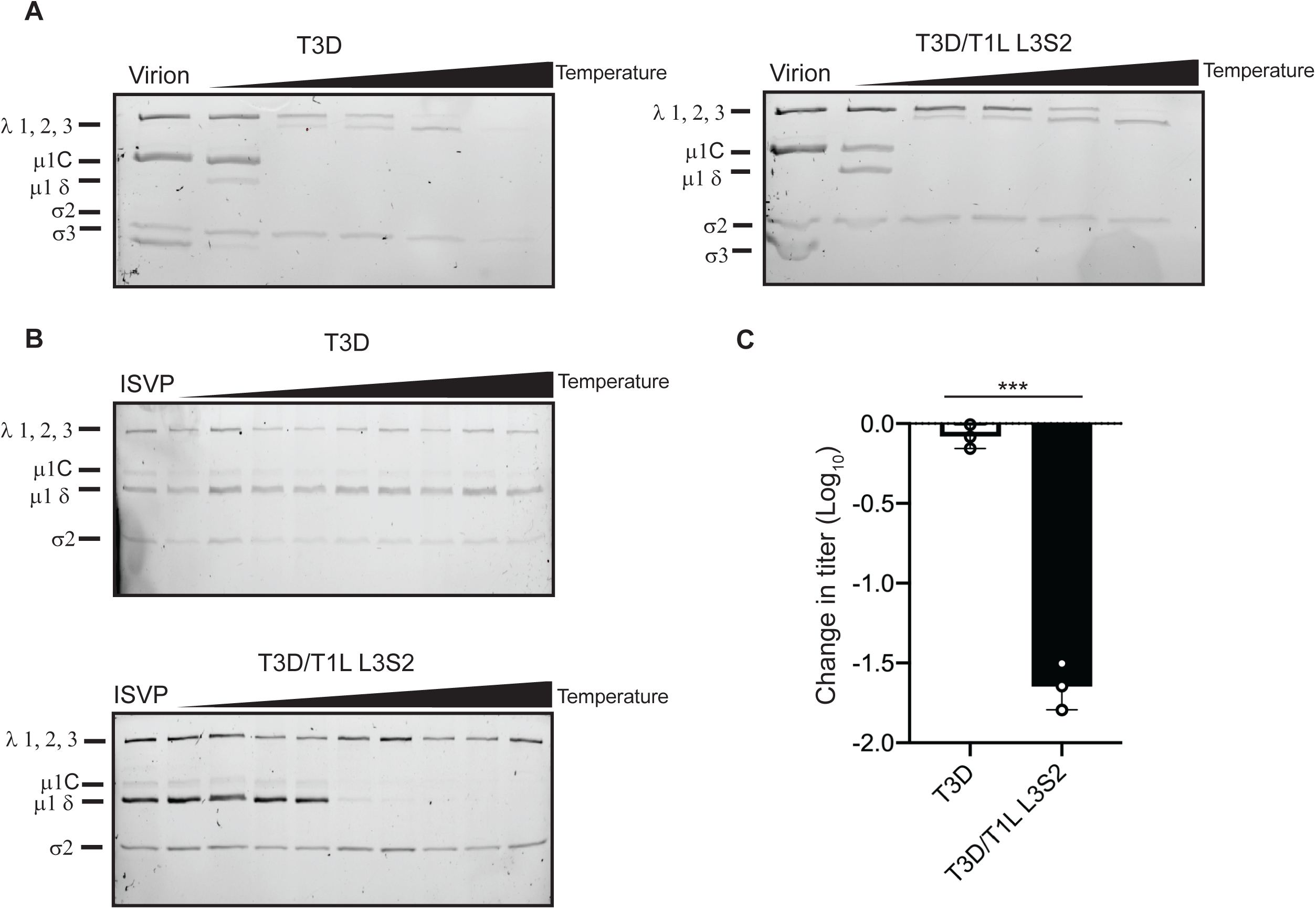
T3D/T1L L3S2 exhibits increased efficiency of ISVP to ISVP* conversion *in vitro*. (A) T3D and T3D/T1L L3S2 virions (2×10^12^ particles/ml) were divided into aliquots of equal volume and incubated at either 4°C or over a range of temperatures (65-85°C) for 5 min. The reactions were chilled on ice and digested with 0.10mg/ml trypsin for 30 min. Following addition of loading dye, the samples were subjected to SDS-PAGE analysis. The positions of major capsid proteins are shown. µ1 runs as µ1C (15). (B) ISVPs (2×10^11^ particles/ml) of T3D or T3D/T1L L3S2 were divided into aliquots of equivalent volume and incubated either at 4°C or over a range of temperatures (22-42°C) for 20 min. The reactions were chilled on ice and digested with 0.10 mg/ml trypsin for 30 min. Following addition of loading dye the samples were subjected to SDS-PAGE analysis. The gels shown are representative of at least 3 independent experiments. The position of major capsid proteins is shown. µ1 runs as µ1C. (C) ISVPs generated from P2 stocks of the indicated virus strain were divided into aliquots of equivalent volume and incubated at either 4°C or 40°C for 20 min. Reactions were then diluted in PBS and subjected to plaque assay. The data are plotted as mean loss of infectivity for three independent samples in comparison to samples incubated at 4°C. Error bars indicate SD. *, P<0.05 as determined by student’s t-test in comparison to T3D.

Conversion to ISVP* results in loss of outer capsid proteins that are essential for entry (27, 28). As a consequence, ISVPs that are heated to temperatures that result in ISVP* conversion have significantly reduced titers. Measuring thermal stability of ISVP infectivity, therefore, is an alternate method to evaluate the efficiency of ISVP* formation. T3D/T1L L3S2 and T3D ISVPs were heated to 40°C and loss of infectivity was measured by plaque assay. Consistent with previous work (20), at 40°C, most of the T3D ISVPs did not convert to ISVP* and the change in titer was minimal (Fig 2C). However, heating T3D/T1L L3S2 ISVPs to 40°C resulted in ISVP* conversion and a significant loss in infectivity (Fig 2C). These data indicate that the reovirus core proteins play a role in stability of the ISVP, a function previously not attributed to the major core proteins λ1 and σ2.

### Enhanced ISVP-to-ISVP* conversion is not due to interactions with RNA

In addition to its structural functions, λ1 plays a variety of other roles during infection. One of those functions involves interaction with RNA (29, 30). If λ1-RNA interactions affect stability of the particles, it may explain why viruses with different core proteins and, consequently, the potential for different RNA interaction properties, may have altered stability. To rule out this possibility, we tested the ISVP-to-ISVP* conversion efficiency of genome-containing particles in comparison to genome-deficient particles of T3D. This comparison would allow us to uncover the contribution of viral genomic RNA to ISVP stability. ISVP-to-ISVP* conversion was again determined as a measure of trypsin sensitivity of µ1 δ. Stability of ISVPs of genome-containing particles and genome-deficient particles was not significantly different which suggests that λ1-RNA interactions do not affect ISVP-to-ISVP* conversion. Thus, the ISVP-to-ISVP* phenotype of T3D/T1L L3S2 is likely not related to differences in RNA interactions (Fig 3).

**Figure 3.**
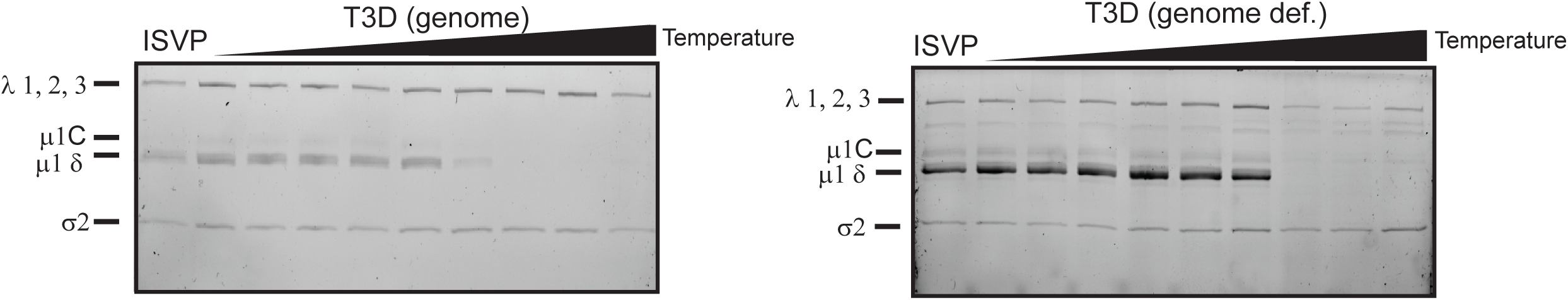
Increased ISVP to ISVP* efficiency in T3D/T1L L3S2 is not due to altered interactions with viral RNA. ISVPs (2×10^11^ particles/ml) derived from genome-containing or genome-deficient particles of strain T3D were divided into aliquots of equivalent volume and incubated at either 4°C or over a range of temperatures (22-40°C) for 20 min. The reactions were chilled on ice and digested with 0.10 mg/ml trypsin for 30 min. Following addition of loading dye the samples were subjected to SDS-PAGE analysis. The position of major capsid proteins is shown. µ1 runs as µ1C.

### Isolation of heat resistant mutants in T3D/T1L L3S2

Because ISVP particles that have converted to ISVP* are less infectious, it is possible to select for and isolate variants with mutations that render ISVPs more stable and thus still retain infectivity after exposure to heat (28, 31, 32). Mapping such mutations could reveal key interactions between viral structural proteins that contribute to maintaining ISVP stability. In order to better understand the basis for the lower stability of this reassortant, we sought to identify such mutations in T3D/T1L L3S2 (Fig 4A). ISVPs of T3D/T1L L3S2 were heated to 40°C (a temperature at which they display significantly lower infectivity than that of similarly treated wild-type ISVPs derived from T3D) and subjected to plaque assay. Resulting plaques were isolated as potential heat resistant mutants. To confirm the heat resistance of these isolates, ISVPs of each isolate were again incubated at 40°C and loss of infectivity in comparison to ISVPs incubated at 4°C was determined by plaque assay. Of the 20 isolates tested, 7 were determined to have little to no loss in infectivity. These isolates were considered to be heat resistant (HR) (Fig 4B). For each of these 7 isolates, genome segments encoding λ1, σ2, and µ1 (L3, S2 and M2 respectively) were sequenced. We reasoned that these genome segments will bear mutations because T3D/T1L L3S2 differs from T3D in the properties of λ1 and σ2, and because µ1 has been previously implicated in controlling stability of ISVPs (28, 33–35). None of the isolates contained mutations in L3 or S2. Four isolates were identified with mutations in µ1. HR16 had a mutation at amino acid 459 (lysine to glutamic acid) which is located in the δ fragment of µ1. HR2, HR15 and HR17 had mutations in the ϕ fragment of µ1. The mutation in HR 15 was at amino acid 607 (proline to glutamine) and HR2 and HR17 had the same mutation at amino acid 615 (alanine to threonine) (Fig 4 C, D, E). Notably, the K459E mutation has been previously identified as a stabilizing mutation (31). However, mutations in µ1 ϕ that contribute to ISVP stability or ISVP-to-ISVP* conversion efficiency have not been previously identified.

**Figure 4.**
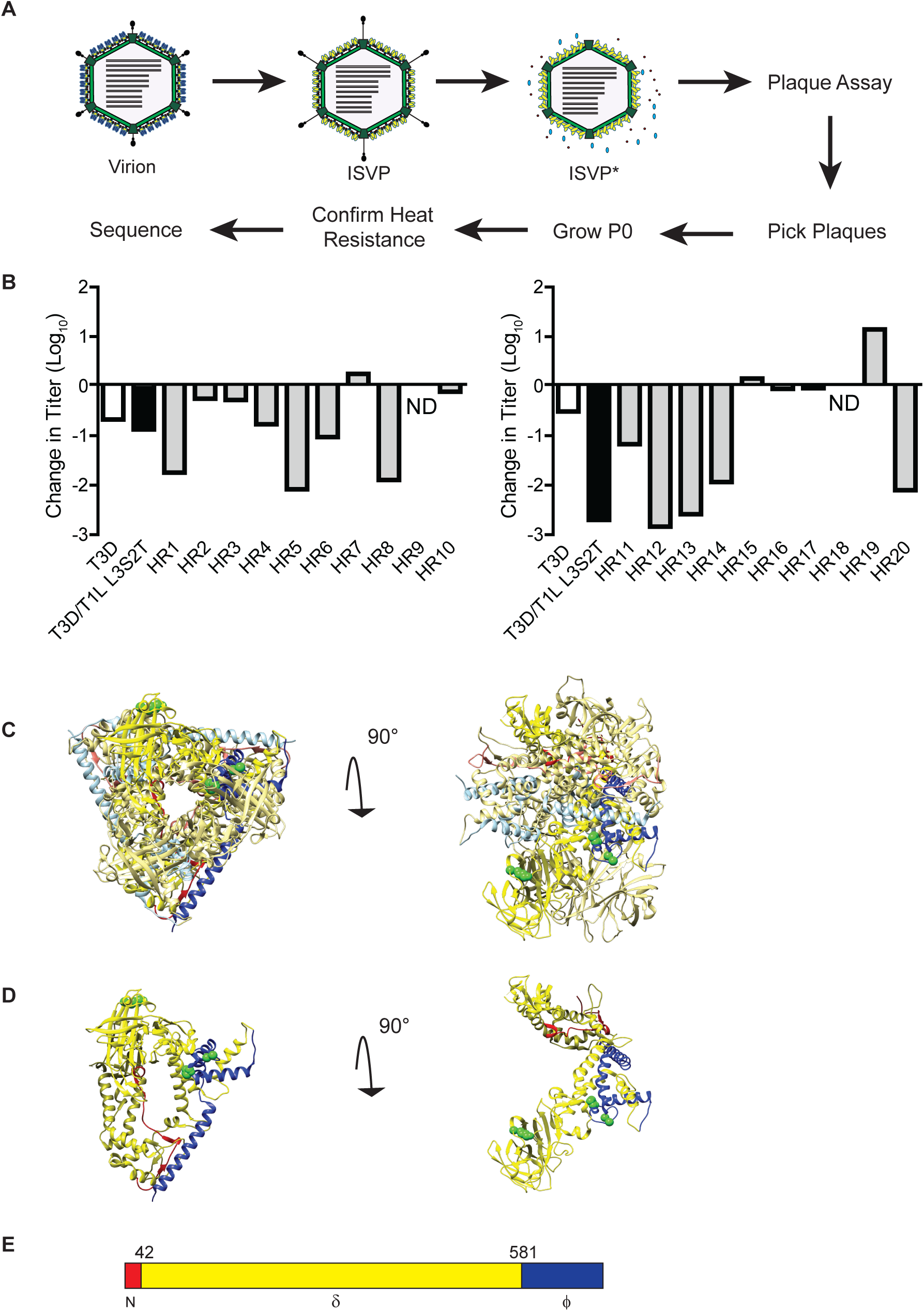
Selection of viruses with mutations that confer stability to T3D/T1L L3S2 ISVPs. (A) Diagram depicting the process for selecting for mutants with reduced ISVP-ISVP* conversion efficiency of T3D/T1L L3S2. ISVPs of T3D/T1L L3S2 were incubated at 40°C for 20 min. Reactions were then diluted in PBS and subjected to plaque assay. Viruses from resulting plaques were isolated and propagated to generate P0 stocks. Heat resistance of these putative heat resistant (HR) plaque isolates was confirmed by measuring the thermal stability of ISVPs incubated at 4°C or 40°C using a plaque assay. Mutants that were confirmed as heat resistant were sequenced. (B) ISVPs generated from P0 stocks were incubated at either 4°C or 40°C for 20 min. Reactions were then diluted in PBS and subjected to plaque assay. ND, Not detectable. (C, D) Top (left) and side (right) views of µ1 trimer (C) and monomer (D) are shown. Position of mutations identified in HR viruses are shown in green. µ1 cleavage fragments are colored as indicated (E) with one µ1 monomer shown with darker colors.

### Mutations in µ1 stabilize T3D/T1L L3S2 ISVPs

Because we did not sequence the entire genome of HR viruses, it remains possible that mutations in genome segments other than L3, S2 and M2 influence the thermal stability of ISVPs generated from the second-site revertants. To evaluate the stabilizing effect of the identified mutations on T3D/T1L L3S2 ISVPs, each mutation was introduced individually into a T3D/T1L L3S2 background. Each new mutant virus was tested for ISVP-to-ISVP* conversion efficiency. ISVPs of each virus were generated and incubated over a gradient of temperatures. ISVP* conversion was determined as a measure of trypsin sensitivity of the µ1 δ fragment (21). In comparison to T3D/T1L L3S2, each of the ISVPs with µ1 mutations had increased stability, suggesting that these amino acid residues in µ1 play important roles in ISVP-to-ISVP* conversion (Fig 5A). To verify these results, the infectivity of ISVPs of T3D/T1L L3S2 and each of the µ1 mutants at 4°C and 40°C was compared by plaque assay. Consistent with the results seen in Fig 2C, T3D/T1L L3S2 ISVPs experience a loss of infectivity at this temperature. In contrast, introduction of µ1 changes identified in heat resistant mutants into T3D/T1L L3S2 result in ISVP particles that display greater stability (Fig 5B). These data indicate that mutations in µ1 are sufficient to restore wild-type like ISVP-to-ISVP* conversion efficiency and thermal stability to T3D/T1L L3S2.

**Figure 5.**
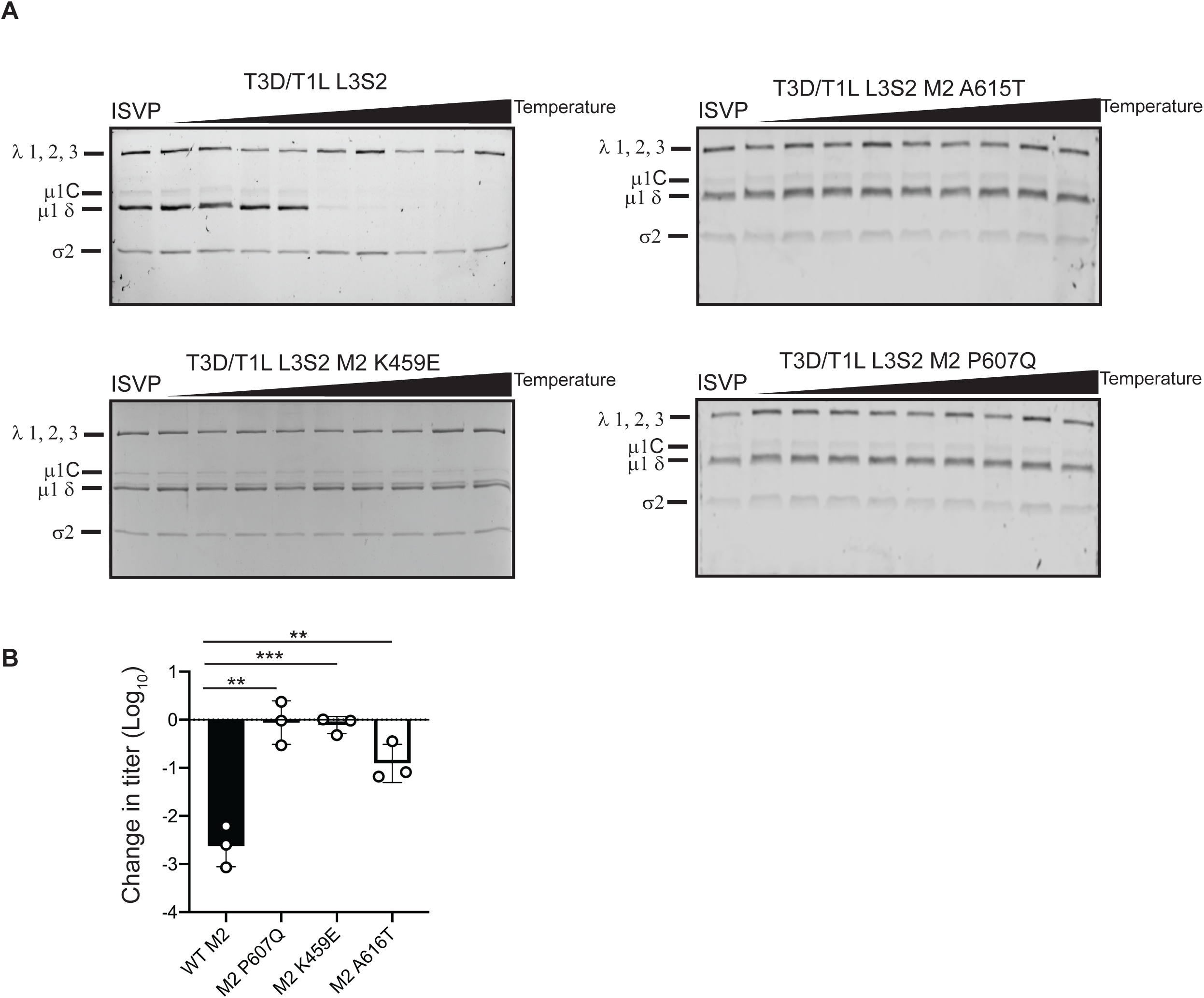
Mutations in µ1 restore stability. (A) ISVPs (2×10^11^ particles/ml) of T3D/T1L L3S2 with the indicated M2 mutations were divided into aliquots of equivalent volume and incubated at either 4°C or over a range of temperatures (22-42°C) for 20 min. The reactions were chilled on ice and digested with 0.10 mg/ml trypsin for 30 min. Following addition of loading dye the samples were subjected to SDS-PAGE analysis. The gels shown are representative of at least 3 independent experiments. The position of major capsid proteins is shown. µ1 runs as µ1C. (B) ISVPs generated from P2 stocks of the indicated virus strain were divided into aliquots of equivalent volume and incubated at either 4°C or 40°C for 20 min. Reactions were then diluted in PBS and subjected to plaque assay. The data are plotted as mean loss of infectivity for three independent samples in comparison to samples incubated at 4°C. Error bars indicate SD. **, P<0.01, ***, P<0.001 as determined by student’s t-test in comparison to T3D/T1L L3S2.

### Mutations in µ1 also affect ISVP-to-ISVP* conversion in wild-type T3D

The µ1 mutations that restored thermal stability of ISVPs and normal ISVP-to-ISVP* conversion efficiency of T3D/T1L L3S2 are not in a position to contact proteins that make up the core (25, 36). Thus, it seems unlikely that these mutations stabilize the capsid by directly altering core-outer capsid interactions. One possibility is that the changes in µ1 simply stabilize the capsid by strengthening interactions between µ1 monomers or between µ1 trimers. If so, the µ1 mutations would be expected to further stabilize ISVPs of T3D, which contains different L3 and S2 alleles. To test this idea, we also generated viruses containing one of each of the three µ1 changes found in the HR viruses in a wild-type T3D background. As before, ISVPs of each virus were generated and incubated over a gradient of temperatures. ISVP* conversion was again determined as a measure of trypsin sensitivity of the µ1 δ fragment. In comparison to wild-type T3D, each mutant underwent ISVP-to-ISVP* conversion much less efficiently (Fig 6A). To verify these results, each virus was tested for loss of infectivity. ISVPs of T3D along with each mutant were heated to 49°C and loss in infectivity was measured by plaque assay. The 49°C temperature was determined empirically as the lowest temperature at which T3D ISVPs exhibit a loss in infectivity (data not shown). Consistent with the data in Fig 6A, at this temperature each of the µ1 mutants displayed no loss of infectivity when compared to wild-type T3D. These data suggest that the mutations identified in µ1 are generally stabilizing mutations and not directly related to the effects of the mismatched core proteins.

**Figure 6.**
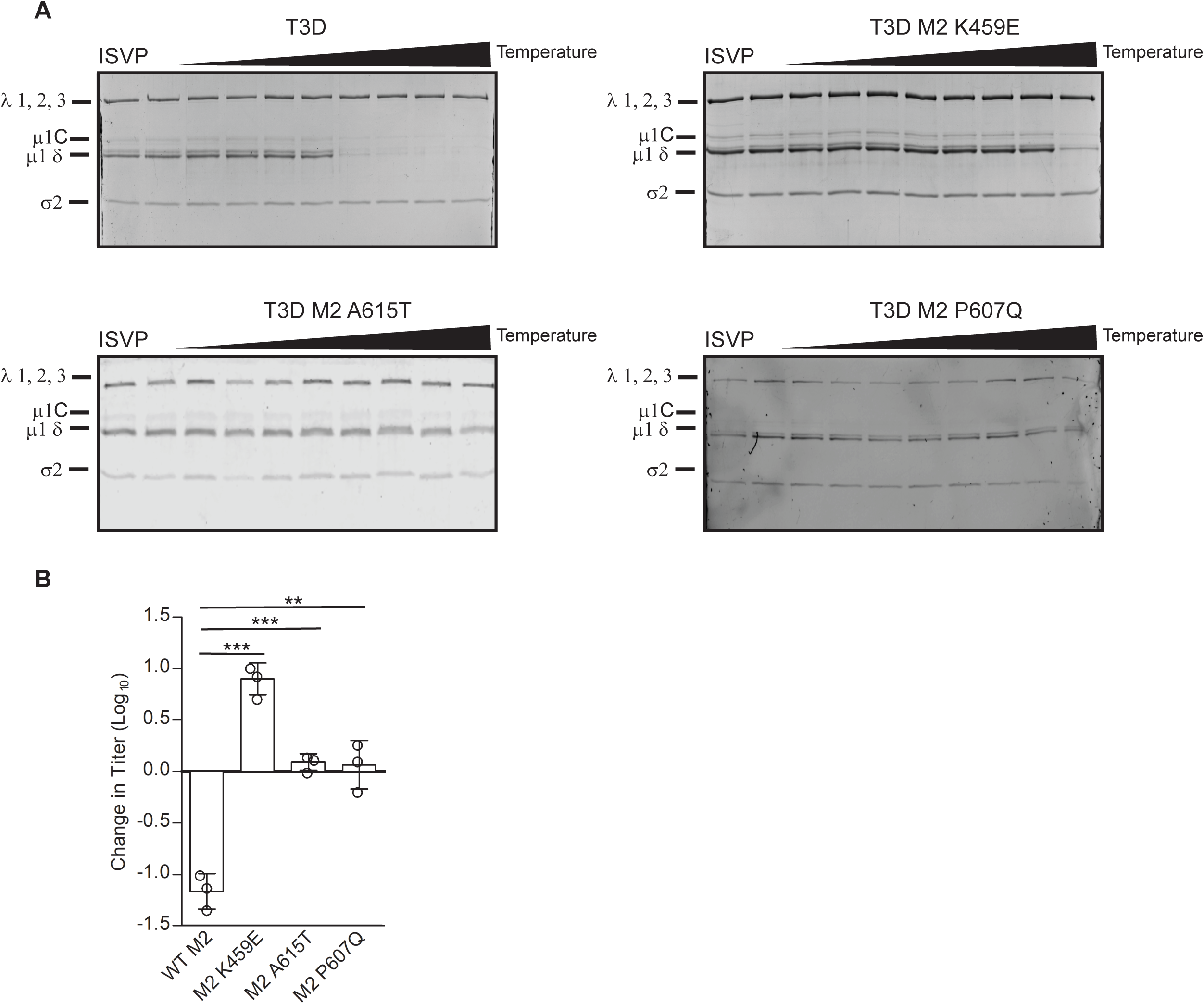
Mutations in µ1 hyperstabilize T3D. (A) ISVPs (2×10^11^ particles/ml) of T3D and T3D with the indicated M2 mutations were divided into aliquots of equivalent volume and incubated at either 4°C or over a range of temperatures (32-46°C) for 20 min. The reactions were chilled on ice and digested with 0.10 mg/ml trypsin for 30 min. Following addition of loading dye the samples were subjected to SDS-PAGE analysis. The gels shown are representative of at least 3 independent experiments. The position of major capsid proteins is shown. µ1 runs as µ1C. (B) ISVPs generated from purified virions were divided into aliquots of equivalent volume and incubated at either 4°C or 42°C for 20 min. Reactions were then diluted in PBS and subjected to plaque assay. The data are plotted as mean loss of infectivity for three independent samples in comparison to samples incubated at 4°C. Error bars indicate SD. ***, P<0.001, **, P<0.01 as determined by student’s T-test in comparison to T3D.

### Mismatches between the core and outer capsid proteins affect viral replication and transcription

To determine if the differences in the ISVP-to-ISVP* conversion efficiency of T3D and T3D/T1L L3S2 are relevant during a viral infection, we next examined viral growth via following infection of cells at an MOI of 0.1 PFU/cell. Virus titer at 24 h following infection was determined by plaque assay and viral yield was calculated as an increase in titer from 0 h post infection (which measured virus adsorbed to cells at the start of infection). We observed that infection with T3D/T1L L3S2 resulted in an increased yield in comparison to T3D (Fig 7A). To determine if this growth phenotype correlates with increased ISVP-to-ISVP* conversion efficiency, we tested the impact of introducing a representative µ1 mutation, A615T in these viruses. While growth of T3D was significantly higher than growth of T3D M2 A615T, growth of T3D/T1L L3S2 was not significantly different from T3D/T1L L3S2 M2A615T (Fig 7A). As the M2 A615T mutation in T3D/T1L L3S2 restores the stability of its ISVPs, these data indicate that differences in replication efficiency of T3D and T3D/T1L L3S2 is not a consequence of the capacity of the reassortant virus to more easily convert to ISVP*. These data also suggest that T1L derived L3 and S2 genome segments can influence the properties of T3D in multiple independent ways.

**Figure 7.**
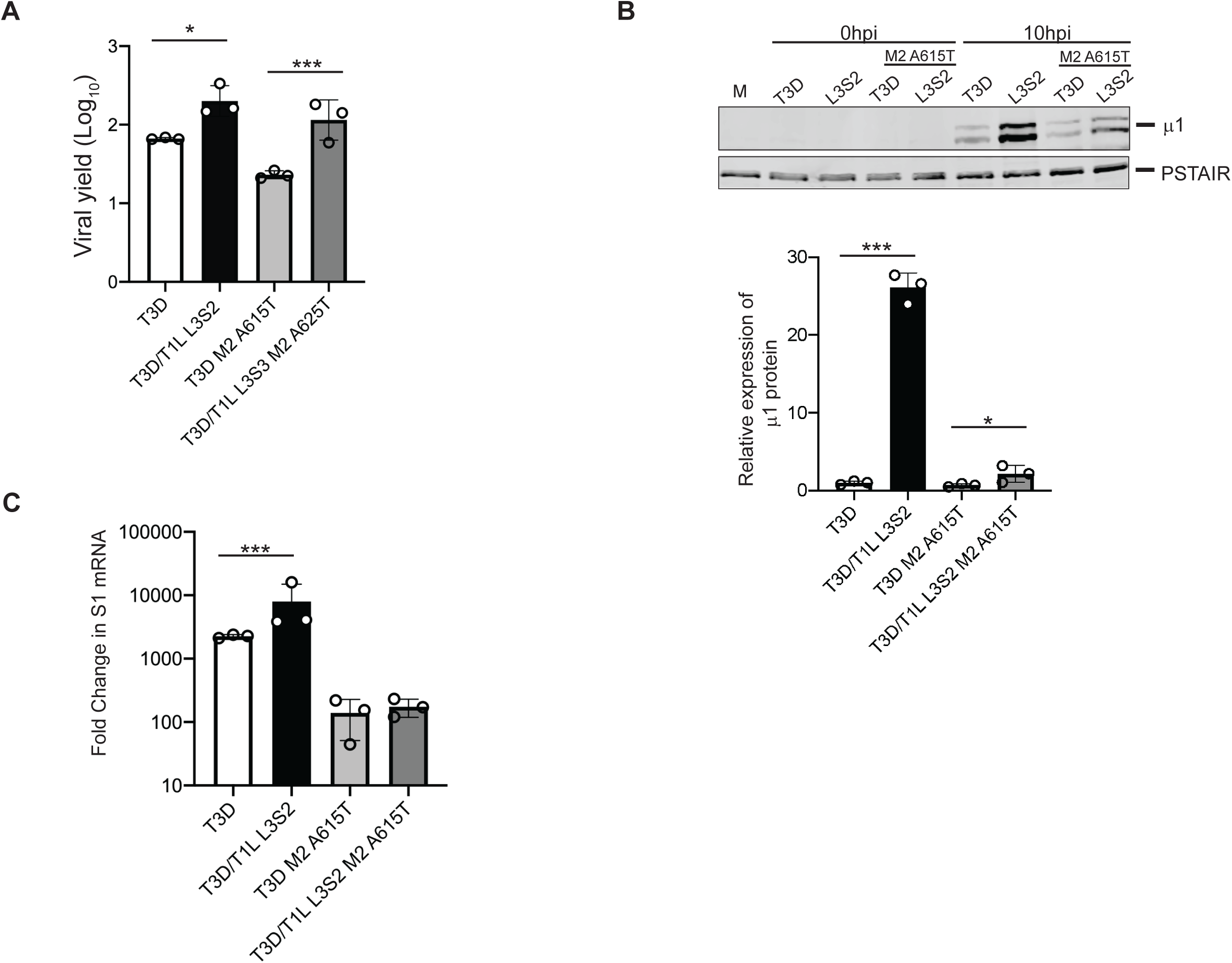
T3D/T1L L3S2 affects viral replication. (A) L cell monolayers were infected with T3D or T3D/T1L L3S2 or with the indicated mutant viruses at an MOI of 0.1 PFU/cell. At 0h and 24h post infection, the infected cells were lysed and the viral yield was quantified by plaque assay. Error bars indicate SD. *, P<0.05, ***, P<0.001 as determined by student’s t-test in comparison to T3D. (B) L cell monolayers were infected with the indicated viruses at an MOI of 10 PFU/cell. At 10h post infection, the cells were lysed and protein production was determined by immunoblotting. Protein quantification of 3 replicates normalized to PSTAIR is shown. Error bars indicate SD. **, P<0.01, ***, P<0.001 as determined by student’s t-test in comparison to T3D. (C) L cell monolayers were infected with the indicated viruses at an MOI of 10 PFU/cell. At the indicated times post infection, the cells were lysed and total RNA was isolated. cDNA was generated using primers for T3D S1 and GAPDH. mRNA production was measured by qPCR. Data shown are represented as fold change compared to mock infected samples and normalized to GAPDH. Error bars indicate SD. ***, P<0.001 as determined by student’s t-test in comparison to T3D

The higher replication potential of T3D/T1L L3S2 could be related to the capacity of this virus to produce more viral gene products with faster kinetics or to a greater extent. To test this idea, we next examined viral gene expression early during infection using immunoblots. Cells were infected with ISVPs at an equal MOI and harvested at 0 or 10h post infection. Expression of the µ1 protein was assessed as a representative. While expression of μ1 was visible at 10 h post infection in all samples, T3D/T1L L3S2 displayed significantly higher protein expression than WT T3D (Fig 7B). To test the impact of ISVP-to-ISVP* conversion phenotypes on protein expression, we also examined T3D M2 A615T and T3D/T1L L3S2 M2 A615T. While both viruses displayed lower protein levels than T3D/T1L L3S2, the mutant in the T3D background had significantly lower μ1 expression than the mutant in the T3D/T1L L3S2 suggesting that the ISVP-to-ISVP* conversion phenotype alone is not responsible for the increase in protein expression. To determine whether the greater level of viral protein expression is a consequence of a higher level of viral mRNA, we measured transcription of viral S1 mRNA early during infection using RT-qPCR. Cells were infected with ISVPs at an equal MOI and harvested at 6 hours post infection. At this timepoint, T3D/T1L L3S2 showed significantly greater reovirus S1 transcripts compared to T3D, indicating that the increase in protein production is likely due to enhanced transcription by the reassortant virus (Fig 7C). These data indicate that swapping core proteins between T1L and T3D in a reassortant virus also influence post-entry viral replication events.

## DISCUSSION

To test the role of the inner core proteins on disassembly and early entry events we generated a reassortant virus with major core proteins from T1L and all other proteins (including the outer capsid) from T3D (T3D/T1L L3S2). T3D/T1L L3S2 undergoes ISVP-to-ISVP* conversion much more efficiently. We identified mutations in µ1 that stabilize ISVPs. T3D/T1L L3S2 also demonstrated increased growth kinetics compared to the parental T3D strain. Surprisingly, the more efficient growth of T3D/T1L L3S2 was not related to its capacity to undergo more efficient ISVP-to-ISVP* transition. Instead, increased growth of T3D/T1L L3S2 relates to more rapid protein and mRNA production early in infection. These data suggest that alteration in properties of core proteins can impact the function of the outer capsid proteins in cell entry. Additionally, properties of core proteins can also influence the enzymatic functions of the capsid that are required to establish efficient infection of host cells.

The reovirus core is a T=1 icosahedron and it is surrounded by the outer capsid (T=13). The outer capsid is made up of 200 heterohexamers of µ1 and σ3 proteins that cover the core (25, 36). It is penetrated by λ2 pentameric turrets at each five-fold axis of symmetry. Trimers of the σ1 attachment protein are situated inside these λ2 turrets (8, 25). The core is made up of 120 copies of λ1 arranged in asymmetric pairs of pentamers to form decamers. Twelve such decamers make up the core. The σ2 protein clamps onto λ1 at 3 different sites within an asymmetric unit resulting in 150 copies of σ2 stabilizing the core shell (8). At each five-fold axis of symmetry, channels penetrate the shell via the λ2 pentameric turrets. It is at each of these channels that the λ3 polymerase interacts with λ1 and is thought to interact with the polymerase co-factor µ2 (5, 37, 38). There are no known contacts between λ1 and the outer capsid. A majority of contacts between the core and the outer capsid occur between µ1 and σ2. These interactions involve the bottom surface of µ1 and the top surface of σ2 (25). The “hub and spoke” structure formed by the C-terminal 33 residue s of µ1 which is thought to be important for stabilizing the µ1 lattice also makes contact with λ2 and/or σ2 (25). This interaction is thought to be stabilized in part by the µ1 residues 51-62 which form flexible loops (25). It is possible that the polymorphic differences between σ2 proteins of T1L and T3D influence interaction with µ1 sufficiently enough such that disassembly is altered. λ1 can also impact interaction of the core with µ1, thereby changing disassembly. However, because µ1 does not contact λ1, this effect would occur indirectly if the structure or conformation of λ2 or σ2, two proteins that do interact with µ1, is altered due to differences in λ1 residues. The relative contribution of differences in properties of σ2 and λ1and the potential subtle differences in structure remain the focus of our ongoing work.

Differences in entry efficiency between different serotypes and laboratory strains of reovirus along with genetic approaches have been used to study how conformational changes and cleavage events required for entry are regulated. Panels of reassortant viruses link differences in ISVP-to-ISVP* conversion to µ1 (28). The autocatalytic cleavage of µ1 and efficiency of ISVP* formation have both been linked to distinct portions of the δ fragment of µ1 (33). Multiple other studies have linked efficiency of entry related disassembly events to the δ fragment of µ1 (31, 33, 35, 36, 39). The µ1N fragment is released from the particle during ISVP* conversion and the released fragment is thought to function in a positive feedback loop to further drive ISVP* conversion (40). The ISVP* promoting activity of µ1N is most efficient in presence of membranes, likely because membrane associated µ1N recruits ISVP-like particles (41, 42). Cleavage of ϕ, which is required for its release from particles during ISVP* conversion is also required for efficient interaction of ISVPs with membranes (42). However, precisely how ϕ functions in this step is not known. Here, we have identified two mutations in the ϕ fragment of µ1 that influence ISVP-to-ISVP* conversion. These mutations are the first indication that the ϕ fragment may also be involved in ISVP-to-ISVP* conversion efficiency. Because our ISVP-to-ISVP* reactions were performed in absence of membranes, the mutations are unlikely to influence ISVP-to-ISVP* conversion by affecting particle-membrane interaction. Thus, the precise mechanism by which the identified ϕ residues influence ISVP* remains unclear. While the majority of µ1-σ3 interactions occur in the jelly-roll domains (residues 306-514) of µ1, both of the mutations identified are in a region proposed to form the cradle for the base of σ3 (36) This region has not been shown to interact with any core proteins or with neighboring µ1 monomers (36). Therefore, it is unlikely that these mutations stabilize ISVPs by strengthening inter- or intra-µ1 trimer interactions or by interacting with core proteins. The identified mutations are in a region of µ1 that may unfold in order to accommodate µ1N release which is a necessary step of ISVP-to-ISVP* conversion (25, 36). Thus, one likely explanation is that ϕ properties affect the release of µ1N.

The L3 gene segment encoded λ1 protein is thought of first as a structural protein that makes up the inner core of reovirus. However, as is the case with most viruses, proteins are capable of playing multiple roles during infection. In addition to its structural roles λ1 has RNA helicase activity, phosphohydrolase activity and is known to interact with RNA (30). The role of each of these functions during infection is currently unknown. The polymerase λ3 interacts with λ1 on the inside of the shell at each five-fold axis (8, 43). While the position of the transcription cofactor protein µ2 within the capsid is not known, it is possible that it also interacts with λ1. Thus, the interaction of λ1 with these encapsidated enzymes could alter viral transcription efficiency. Reovirus serotypes T1L and T3D replicate with different efficiency in some cell lines, with T1L replicating to a higher extent and with faster kinetics (44). Reassortant analyses have partially linked this phenotype to the T1L derived λ1-encoding L3 gene segment (44). Our studies indicate that the enhanced infection efficiency of T3D/T1L L3S2 is not related to its greater propensity for ISVP-to-ISVP* transition. Instead, we propose that enhanced efficiency of infection is a result of differences in the activity of λ1 itself or its impact on the activity of the transcriptional machinery comprised of λ3 and µ2. While the role of σ2 as a structural protein is well established, additional roles in replication have not been confidently identified. It has weak interactions with dsRNA and reassortant studies have linked σ2 with increased induction and sensitivity to interferon in some cell types (45, 46). Additional studies are needed with monoreassortants bearing only S2 and L3 gene segments in the T3D background to precisely ascertain the basis for the enhanced replicative efficiency of T3D/T1L L3S2.

Recent studies from our laboratory have revealed that reassortant viruses display phenotypes that are unexpected and extend beyond the known function of the protein. First, we found that, even though the primary function of the M2 encoded protein µ1 is in membrane penetration, an M2 reassortant virus displays greater attachment to host cells (47). Second, we found that though the S1 encoded σ1 protein is the attachment factor, an S1 reassortant impacts the stability of the µ1 layer with which it makes no interactions (20). Our current study presented here reveals that core proteins, previously only thought to have a structural role in packaging the genomic material, influence cell entry events regulated by the outer capsid. Until the advent of reverse genetics, reassortant analyses have been used as the main strategy to assign function to proteins of segmented viruses. This approach has also been useful to determine the genetic basis of viral disease. While this approach has been invaluable, we think our work suggests that the structure-function explanation of some phenotypes reported for reovirus and possibly other members of the *Reoviridae* family may be more complicated than previously appreciated.

## MATERIALS AND METHODS

### Cells and viruses

Spinner adapted murine L929 (L) cells were grown at 37°C in Joklik’s minimal essential medium (Lonza) supplemented with 5% fetal bovine serum (FBS) (Life Technologies), 2 mM L-glutamine (Invitrogen), 100 U/ml penicillin (Invitrogen), 100 μg/ml streptomycin (Invitrogen), and 25 ng/ml amphotericin B (Sigma-Aldrich). All virus strains used in this study were derived from reovirus type 3 Dearing (T3D) and reovirus type 1 Lang (T1L) and were generated by plasmid-based reverse genetics (48). Mutations within the T3D M2 gene were generated by QuikChange site-directed mutagenesis (Agilent Technologies). Primer sequences are available upon request.

### Virus propagation and purification

All wild-type and mutant viruses used in this study were propagated and purified as previously described (49, 50). Briefly, plaques isolated from plasmid based reverse genetics were propagated successively in T-25, T75 and T-175 flasks to generate P0, P1 and P2 virus stocks respectively. To generate purified virus, L cells infected with P2 reovirus stocks were lysed by sonication. Virus particles were extracted from the lysates using Vertrel-XF specialty fluid (Dupont) (51). The extracted particles were layered onto 1.2- to 1.4-g/cm^3^ CsCl step gradients. The gradients were then centrifuged at 187,000 × *g* for 4 h at 4°C. Bands corresponding to purified virus particles (∼1.36 g/cm^3^) (52) were isolated and dialyzed into virus storage buffer (10 mM Tris, pH 7.4, 15 mM MgCl_2_, and 150 mM NaCl). Following dialysis, the particle concentration was determined by measuring the optical density of the purified virus stocks at 260 nm (OD_260_) (1 unit at OD_260_ is equal to 2.1 × 10^12^ particles/ml)

### Generation of ISVPs

Purified virions of the indicated virus strains (2 × 10^12^ particles/ml or 4 × 10^12^ particles/ml) were digested with 200 μg/ml TLCK (*N*α-*p*-tosyl-L-lysine chloromethyl ketone)-treated chymotrypsin (Worthington Biochemical) in a total volume of 100 μl for 1 hour at 32°C. After 1 h, the reaction mixtures were incubated for 20 min on ice and quenched by the addition of 1 mM phenylmethylsulfonyl fluoride (Sigma-Aldrich). The generation of ISVPs was confirmed by SDS-PAGE and Coomassie brilliant blue staining.

### Analysis of ISVP-ISVP* conversion

ISVPs (2 × 10^12^ particles/ml) of the indicated viral strains were divided into aliquots of equivalent volumes and heated at the indicated temperatures for 20 min. The reaction mixtures were cooled on ice and then digested with 0.10 mg/ml trypsin (Sigma-Aldrich) for 30 min on ice. Following addition of the SDS-PAGE loading dye, the samples subjected to SDS-PAGE analysis. For analysis by quantitative infectivity assay, P2 stocks or purified virus stocks of the indicated viruses were diluted 1:10 in virion storage buffer (10 mM Tris, pH 7.4, 15 mM MgCl_2_, and 150 mM NaCl). 200µg/mL TLCK (*N*α-*p*-tosyl-L-lysine chloromethyl ketone)-treated chymotrypsin (Worthington Biochemical) was added to each sample. Samples were heated to 37°C for 30 min. The reaction was quenched by the addition of 1 mM phenylmethylsulfonyl fluoride (Sigma-Aldrich) and cooled on ice for 10 min. The reactions were divided in equivalent volumes and incubated at 4°C or 40°C for 20 min. Reactions were used to initiate infection of L929 cells and infectivity was determined by plaque assay. The change in infectivity at a given temperature (*T*) was calculated using the following formula: log_10_(PFU/ml)_*T*_ − log_10_(PFU/ml)4°C.

### Analysis of virion stability

Virions (2×10^12^ particles/ml) of the indicated viral strains were divided into aliquots of equivalent volumes and heated at the indicated temperatures for 20 min. The reaction mixtures were cooled on ice and then digested with 0.10 mg/ml trypsin (Sigma-Aldrich) for 30 min on ice. Following addition of SDS loading dye, the samples were subjected to analysis by SDS-PAGE.

### Isolation and verification of heat resistant (HR) mutants

ISVPs of purified T3D/T1L L3S2 were generated and subsequently heated to 40°C for 20 min. Resulting reactions were diluted in phosphate-buffered saline (PBS) supplemented with 2 mM MgCl_2_ and subjected to plaque assay. Heat resistant mutants were selected by plaque purification and propagated in L cells to obtain P0 viral stocks. P0 stocks were diluted 1:10 in virion storage buffer (10 mM Tris, pH 7.4, 15 mM MgCl_2_, and 150 mM NaCl). 200µg/mL TLCK (*N*α-*p*-tosyl-L-lysine chloromethyl ketone)-treated chymotrypsin (Worthington Biochemical, Lakewood, NJ) was added to each sample. Samples were heated to 37°C for 30 min. The reaction was quenched by the addition of 1 mM phenylmethylsulfonyl fluoride (Sigma-Aldrich) and cooled on ice for 10 min. The reactions were divided in equivalent volumes and incubated at 4°C or 40°C for 20 min. Reactions were used to initiate infection of L929 cells and infectivity was determined by plaque assay. The change in infectivity at a given temperature (*T*) was calculated using the following formula: log_10_(PFU/ml)_*T*_ − log_10_(PFU/ml)4°C.

### Plaque titration

Plaque assays were conducted in spinner-adapted L929 cells plated in 6-well plates (Greiner Bio-One). Cells were adsorbed with dilutions of virus in phosphate-buffered saline (PBS). Cells were overlaid with a molten mixture comprised of 1× medium 199 and 1% Bacto agar supplemented with 10 μg/ml chymotrypsin. Five days following infection, the monolayers were fixed by addition of 4% formaldehyde solution in PBS and incubated overnight. The agar overlay was peeled off, and the monolayers were stained with 1% crystal violet stain in 5% ethanol for 5 h at room temperature. The monolayers were washed with water. Virus titer was quantified by manual counting of plaques.

### Analysis of protein levels by immunoblotting

The samples were whole-cell lysates of infected cells prepared using radioimmunoprecipitation assay (RIPA) lysis buffer (50 mM NaCl, 1 mM EDTA at pH 8, 50 mM Tris at pH 7.5, 1% Triton X-100, 1% sodium deoxycholate, 0.1% SDS) supplemented with protease inhibitor cocktail (Roche) and 500 μM PMSF, and they were resolved on 10% SDS-PAGE gels and transferred to nitrocellulose membranes. For immunoblotting using polyclonal rabbit antireovirus serum, the membranes were blocked with 5% milk in Tris-buffered saline (TBS) at room temperature for 1 h. Following blocking, rabbit anti-reovirus serum (1:1,000) or anti-PSTAIR was incubated with the membrane in appropriate blocking buffer at room temperature for 1 h. The membranes were washed with TBS supplemented with 0.1% Tween 20 (TBS-T) twice for 15 min and then incubated with Alexa Fluor-conjugated anti-rabbit IgG or anti-mouse IgG in blocking buffer. Following three washes, membranes were scanned using an Odyssey infrared imager (LI-COR), and intensities of bands were quantified using Image Studio Lite software (LI-COR).

### Analysis of mRNA levels by RT-qPCR

RNA was extracted from infected cells, at various times after infection, using a total RNA minikit (Bio-Rad). For RT-qPCR, 0.5 to 2 μg of RNA was reverse transcribed with the high-capacity cDNA RT kit (Applied Biosystems), using random hexamers for amplification of cellular and viral genes. Undiluted cDNA was subjected to PCR using SYBR Select Master Mix (Applied Biosystems) and primers specific for T3D S1 and GAPDH. Fold increases in gene expression with respect to control samples (indicated in figure legend) were measured using the ΔΔ*C_T_* method (53). Calculations for determining ΔΔ*C_T_* values and relative levels of gene expression were performed as follows: fold increase in viral gene expression = 2_[-(ΔΔCT)]_

### Statistical analyses

The reported values represent the means of three independent biological replicates. The error bars indicate standard deviations (SD). *P* values were calculated using Student’s *t* test (two-tailed; unequal variance assumed).

### Modeling

Molecular graphics were created and analysis were performed with the UCSF Chimera package (54).

